# Comparative transcriptomics reveals a conserved Bacterial Adaptive Phage Response (BAPR) to viral predation

**DOI:** 10.1101/248849

**Authors:** Bob G. Blasdel, Pieter-Jan Ceyssens, Anne Chevallereau, Laurent Debarbieux, Rob Lavigne

**Affiliations:** Laboratory of Gene Technology, KU Leuven, Leuven, Belgium; Institut Pasteur, Molecular Biology of the Gene in Extremophiles Unit, Department of Microbiology, Paris, France

**Author notes:** For correspondence (BB); (RL). **Present address:** National Reference Center for Tuberculosis and Mycobacteria, Scienti1c Institute of Public Health (WIV-ISP), Brussels, Belgium.

## Abstract

Intrinsic and acquired defenses against bacteriophages, including Restriction/Modification, CRISPR/Cas, and Toxin/Anti-toxin systems have been intensely studied, with profound scientific impacts. However, adaptive defenses against phage infection analogous to adaptive resistance to antimicrobials have yet to be described. To identify such mechanisms, we applied an RNAseq-based, comparative transcriptomics approach in different *Pseudomonas aeruginosa* strains after independent infection by a set of divergent virulent bacteriophages. A common host-mediated adaptive stress response to phages was identified that includes the Pseudomonas Quinolone Signal, through which infected cells inform their neighbors of infection, and what may be a resistance mechanism that functions by reducing infection vigor. With host transcriptional machinery left intact, we also observe phage-mediated differential expression caused by phage-specific stresses and molecular mechanisms. These responses suggest the presence of a conserved Bacterial Adaptive Phage Response mechanism as a novel type of host defense mechanism, and which may explain transient forms of phage persistence.

## Introduction

*Pseudomonas aeruginosa* is a ubiquitous Gram-negative soil organism which thrives in diverse humid environments and acts as an opportunistic pathogen. Around 8% of its unusually large and complex genome is predicted to encode transcriptional regulators and two-component regulatory systems, allowing it to rapidly adapt to changing conditions (*Stover et al., 2000*). Recently, by analyzing the transcriptomes of the *P. aeruginosa* PA14 type strain under 14 different environmental conditions, *Dötsch et al*. (*2015*) were able to precisely define the parameters and context of *P. aeruginosa* transcription. Indeed, they found that while only 24-46% of gene features are active within each environmental condition that they tested, nearly all gene features were active in at least one.

Bacteriophages (phages) are a dominant force in the control of microbial communities and typically outnumber their bacterial hosts by tenfold. Phage predation accounts for a significant proportion of bacterial mortality, up to 80% in some systems (*Rohwer et al., 2009*), and thus presents a remarkably strong source of selective pressure for bacterial immunity systems. Known bacterial immunity systems can be divided into categories analogous to either intrinsic or acquired antimicrobial resistance systems as denoted by *Arzanlou et al. (2017)*. Indeed, a wide variety of phage defense systems have been found that are analogous to intrinsic mechanisms of antimicrobial resistance, where bacteria constitutively express systems that mediate phage resistance, such as Restriction/Modification systems (*Tock and Dryden, 2005*), Toxin/Anti-toxin systems (*Dy et al., 2014*), and the Bacteriophage Exclusion (BREX) system (*Goldfarb et al., 2015*). Similarly a great diversity of phage immunity systems analogous to acquired antimicrobial resistance systems, where bacteria acquire genetic material that engenders resistance to phages, have been found such as prophage homoimmunity (*Adams et al., 1959*), DNA-guided DNA interference by prokaryotic Argonaute (*Swarts et al., 2014*), and CRISPR/Cas-mediated interference (*Marraffini, 2015*). However, to date, it has been unclear to what extent bacteria might possess immunity mechanisms analogous to adaptive antimicrobial resistance systems, where bacteria would recognize a phage threat and express a stress response to phage predation that would protect either the infected cell or its nearby kin.

As true parasites, phages rely heavily on their host cell mechanisms for their reproduction. When an obligately lytic phage injects its genome into the cytoplasm of a bacterial host, a molecular battle for dominance over the cell ensues as phage mechanisms attempt to convert the host to a viral metabolism conducive to phage production, while the host mechanisms work to exclude or inhibit viral propagation. Lacking the ability to produce a temperate lifecycle, infection by obligately lytic phages typically results in cell death and either successful or unsuccessful propagation of the phage. Many have assumed that one of the primary ways in which phage dominate their hosts during a lytic life cycle is to eliminate host transcription and transcripts, based on the example of T4 (*Kutteret et al., 1994*). However, our recent investigations have shown that *Pseudomonas aeruginosa* phages PAK_P3 and ΦKZ respectively appear to direct the upregulation of an operon involved in host mRNA stability (*Chevallereau et al., 2016*) and prompt the transcriptional mobilization of a prophage (*Ceyssens et al., 2014*). At the same time, *Howard-Varona et al*. (*2016*) have demonstrated that the genomes of two different *Cellulophaga baltica* strains were differentially expressed in unique ways during Φ38:1 phage infection, hinting at the existence of diverse transcription-level stress responses to the one phage examined. However, it has thus far been impossible to tease apart which effects on the stability and expression of host transcripts during infection are phage-mediated and which effects are prompted by the host in response to phage stress. During phage infections where the expression of host genetic information is indeed being meaningfully differentially regulated, it thus becomes imperative to distinguish whether the virus or the host within the cell is directing that differential expression in order to meaningfully understand it.

Here we performed RNA-Seq on independent infections of *P. aeruginosa* by divergent obligately lytic phages and define the host stress response to phage infection by identifying the genes that are differentially expressed in common. This isolates the transcriptional impact of the host mechanisms responding to phage infection that all of these phage infections have in common. Indeed, identifying genes differentially expressed during all of the phage infections will distinguish this from host regulatory responses to stresses imposed by specific viral metabolisms *DeSmet et al. (2016)* as well as the transcriptional impact of phage-specific molecular mechanisms (*Ceyssens et al., 2014*). Our ability to putatively assign the agency behind differential expression to either host or phage has allowed us to describe a plethora of fundamentally novel adaptive host systems for defending against phage predation. This includes the well-studied PQS metabolite, a quorum sensing compound (*Diggle et al., 2003,2007*). *Haussler and Becker (2008)* have previously proposed that the dual nature of PQS, as a pro-oxidant and a regulator with the ability to promote a protective anti-oxidative response, may be involved in rescuing healthy members of the community from oxidative stress while promoting the destruction of unhealthy sister cells whose survival would reduce the fitness of the micro-colony as a whole. However, our results suggest that it also functions as a phage defense warning capable of mediating phage resistance. Our results also demonstrate that, unlike *T4virus* members that have long been demonstrated to shut off host gene expression in immediate-early infection (*Koerner and Snustad, 1979*), many *Pseudomonas* phages maintain the host regulatory apparatus - challenging this long-standing paradigm. Thus we are also able to postulate that, at least some phage manipulate the genetic wealth encoded by the host towards phage ends, exploiting the host genome as if it were an auxiliary phage genome.

## Results and Discussion

### A diverse library of obligately lytic phages

RNA Sequencing was performed on infections of *P. aeruginosa* PAO1 by divergent phages that are completely devoid of genomic synteny or even genome similarity. Samples were collected at time points chosen to represent early, middle, and late infection, using the well established T4 model as a guide (*Miller et al., 2003*), as well as phage negative controls (Samples outlined in Supplementary Table 2). The phages outlined in Table 1 each co-opt host mechanisms in radically different ways and have unique impacts on the abundance of host metabolites (*De Smet et al., 2016*). As a result, they are are unlikely to all mediate differential expression of the same host genes in the same way.

**Table 1.**
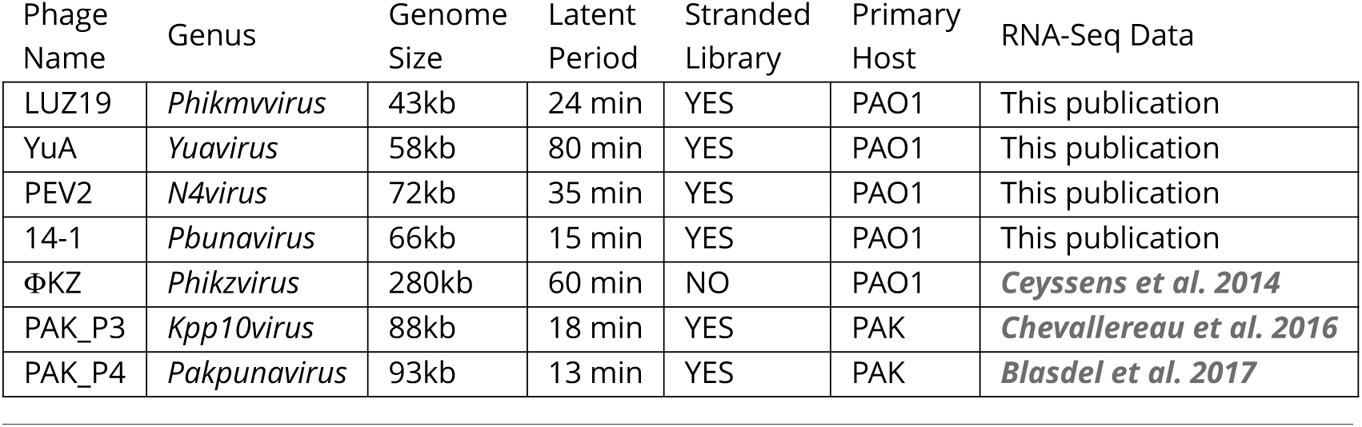
None of the five phage infecting PAO1 described here share any gene products to any other phage, based on sequence-based similarity (*De Smet et al., 2016*), while the relationship of PAK_P3 to PAK_P4 is described by *Blasdel et al*. (*2017*). Data for all phage save OKZ was sequenced from cDNA library preparations that were strand specific, allowing for sense transcripts to be counted independently of antisense transcripts.

| Phage Name | Genus | Genome Size | Latent Period | Stranded Library | Primary Host | RNA-Seq Data |
| --- | --- | --- | --- | --- | --- | --- |
| LUZ19 | *Phikmvirus* | 43kb | 24 min | YES | PAO1 | This publication |
| YuA | *Yuavirus* | 58kb | 80 min | YES | PAO1 | This publication |
| PEV2 | *N4virus* | 72kb | 35 min | YES | PAO1 | This publication |
| 14-1 | *Pbunavirus* | 66kb | 15 min | YES | PAO1 | This publication |
| ΦKZ | *Phikzvirus* | 280kb | 60 min | NO | PAO1 | *Ceyssens et al. 2014* |
| PAK_P3 | *Kpp10virus* | 88kb | 18 min | YES | PAK | *Chevallereau et al. 2016* |
| PAK_P4 | *Pakpunavirus* | 93kb | 13 min | YES | PAK | *Blasdel et al. 2017* |

Within the infections of the seven phages with RNA-Seq data discussed here, there is a great diversity of impacts on the abundance of phage transcripts relative to host transcripts as shown in Figure 1. Indeed, at one extreme, PAK_P4 (*Blasdel et al., 2017*) can be observed to produce 84% of all the transcripts in the cell after 3.5 minutes while at the other end YuA produces less than 5% even until late infection. Some phages such as PAK_P3 appear to replace host transcripts at a constant rate (*Chevallereau et al., 2016*), while others appear to slowly accelerate the replacement such as LUZ19 and ΦKZ, and others such as 14-1 and YuA appear to have a sudden impact at a specific point in the middle of infection. These different impacts appear to instead reflect fundamentally different relationships between the studied phages and the transcriptomes of the host as well as fundamentally different RNA nucleotide metabolisms.

**Figure 1.**
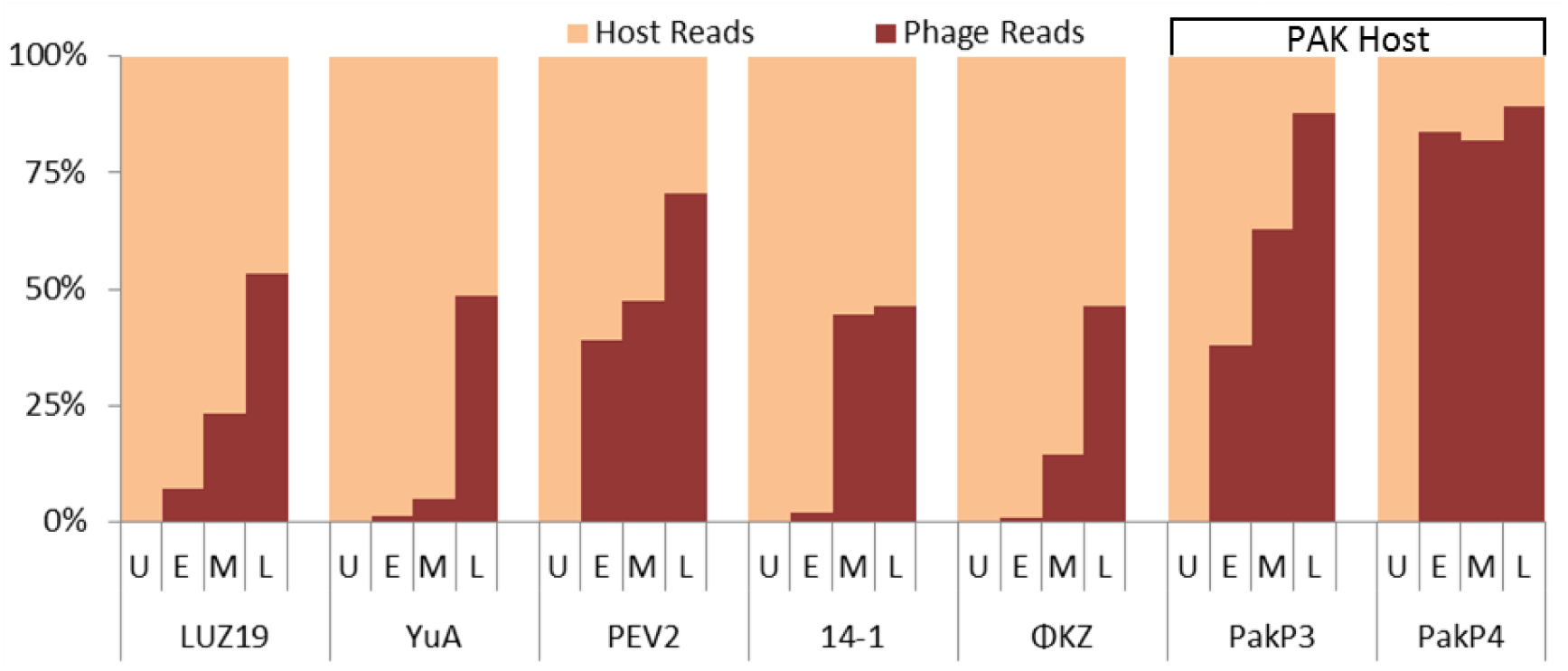
Reads aligning to phage genomes over the course of infection. Shown here is the percentage of reads that map to the phage genome when compared to the total non-ribosomal RNA reads that map to either the phage or host genome. This proportion ends up representing the relative mass of non-ribosomal RNA present in the cell. Also shown for comparison are reads harvested from previously published data for ΦKZ, PAK_P3 (*Chevallereau et al., 2016*), and PAK_P4 (*Blasdel et al., 2017*).

### The putative PAO1 stress response to phage distinguished from phage-mediated impacts

To define the putative host-mediated *P. aeruginosa* PAO1 stress response to phage infection, we compared strand-specific RNA-Seq data representing independent infections by four divergent phages (LUZ19, YuA, PEV2 and 14-1) in the core analysis (See Figure 2), identifying gene features that were differentially expressed in common. This possible stress response to phage infection, which differentially expresses 5.5% of the coding sequences present in *P. aeruginosa* PAO1, contains a number of systems that appear to be involved in hindering phage infection. Indeed, these include an indication of increased Hfq activity through the depletion of the CrcZ sRNA (See Appendix Box 1), a quorum sensing-based phage defense system, and a form of reduced infection vigor. Additional features are highlighted in Appendix 1. However, this putative host-mediated stress response is notably absent during infection by ΦKZ (*Ceyssens et al., 2014*) while the transcripts that are degraded during other infections persist.

**Figure 2.**
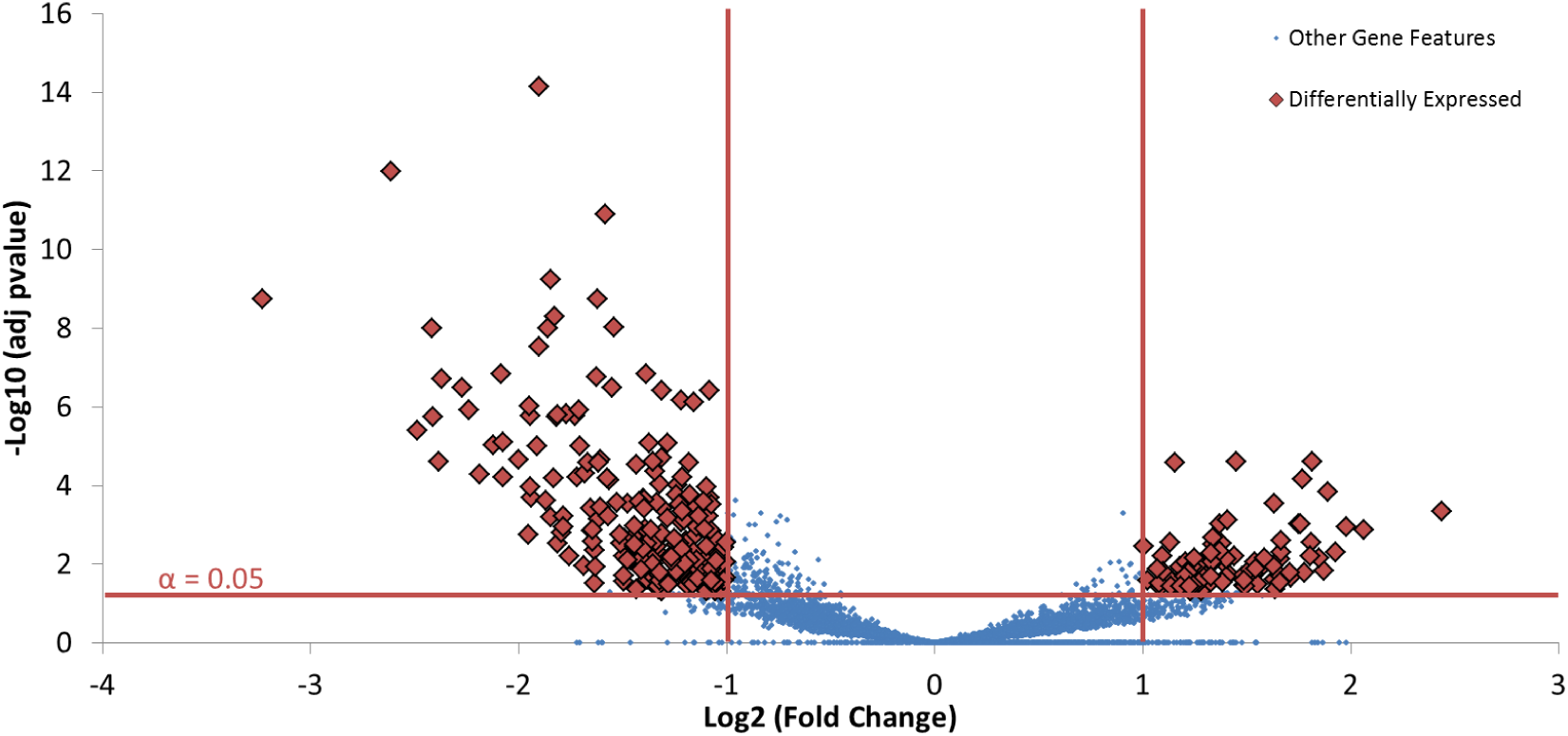
Defining the host response. This volcano plot shows gene features that can be considered differentially expressed (Log2 Fold Change >1 or <-1, and adj pvalue < 0.05) between the phage negative control and the late infections of phages LUZ19, YuA, 14-1, and PEV2 considered as a single condition. This highlights a putative host-mediated stress response to phage infection that is independent of phage-mediated manipulation of host gene expression.

The Pseudomonas Quinolone System is upregulated in response to phage infection One of the most remarkable components of the host-mediated stress response to phage infection is the transcriptional upregulation of the pqsABCDE operon, as well as the co-regulated phnAB operon, which are both necessary for the formation of the Pseudomonas Quinolone Signal (PQS) (*Gallagher et al., 2002*; *Palmer et al., 2013*). As one of the two primary functions of the PQS metabolite appears to be as a transcriptional regulator, upregulating stationary phase expression and the oxidative stress response, we compared our data to the differential expression of Coding Sequences found by *Rampioni et al. (2010)* to be regulated by PQS (See Figure 4). Our results suggest that rather than serve as just a warning of oxidative stress (*Häussler and Becker, 2008*), the PQS family of compounds could also function as a warning of coming phage infection. This would help explain the results of *Moreau et al*. (*2017*), who found that a PQS quorum-signaling proficient strain of *P. aeruginosa* was able to evolve higher levels of resistance to phages during a short-term selection experiment. Indeed, the PQS quorum sensing system could work as a host defense against phages by shutting down the metabolisms of not yet infected cells and weakening or killing phage-infected cells that lack the ability to transcriptionally respond to its toxic effect. While neither effect could plausibly save a cell that has received the chemical signal with any reasonable frequency, both could substantially reduce burst sizes or prevent productive infection, impacting the success of the phage and its propagation within the host community.

**Figure 3.**
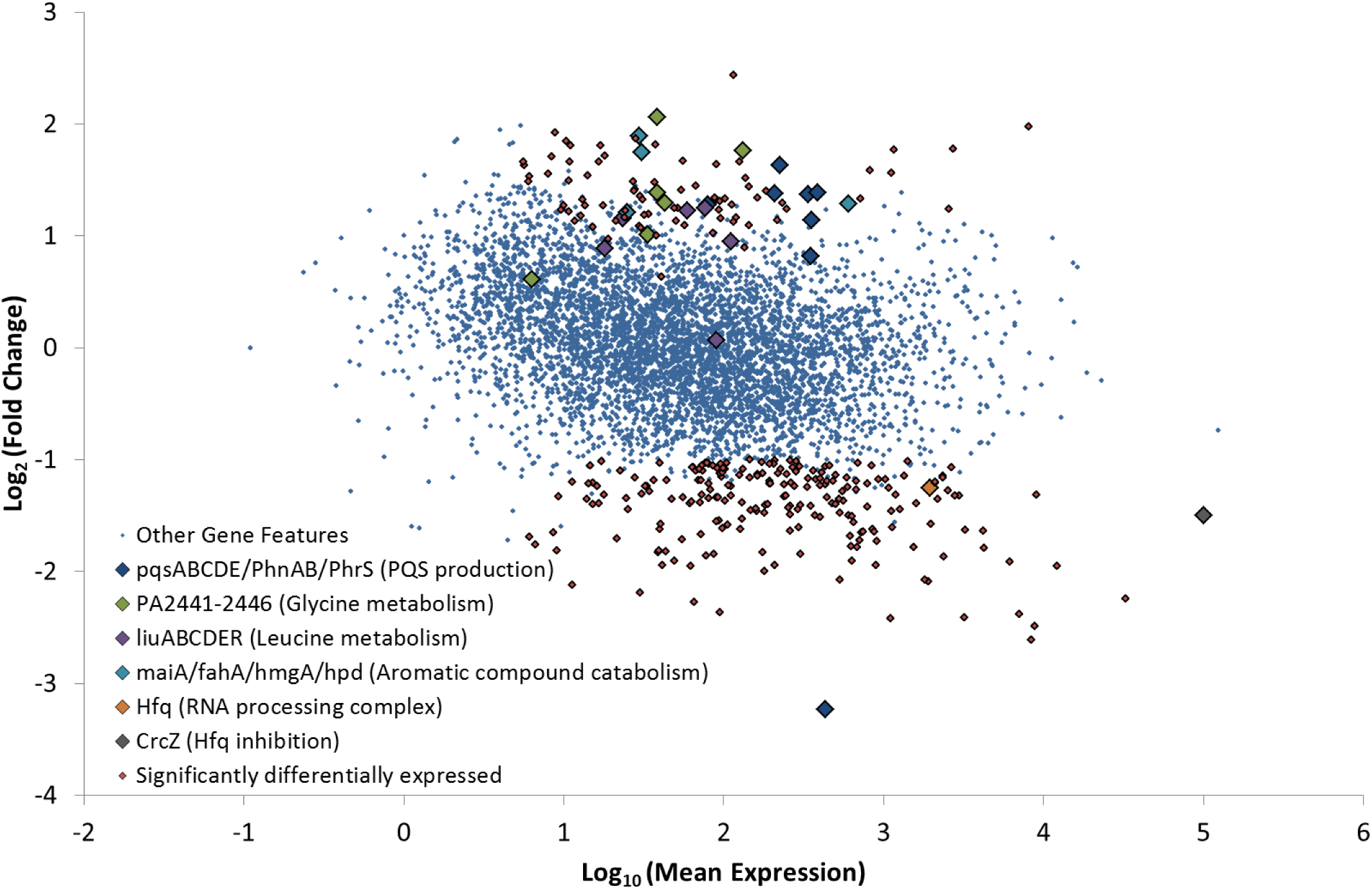
MA Plot highlighting the operons that are differentially expressed in common between infections. By considering samples representing the abundance of host transcripts during the late infections of phages LUZ19, YuA, PEV2 and 14-1 as a single condition and comparing it to a condition representing the phage-negative control, we are able to describe genes that are differentially expressed in common between infections.

**Figure 4.**
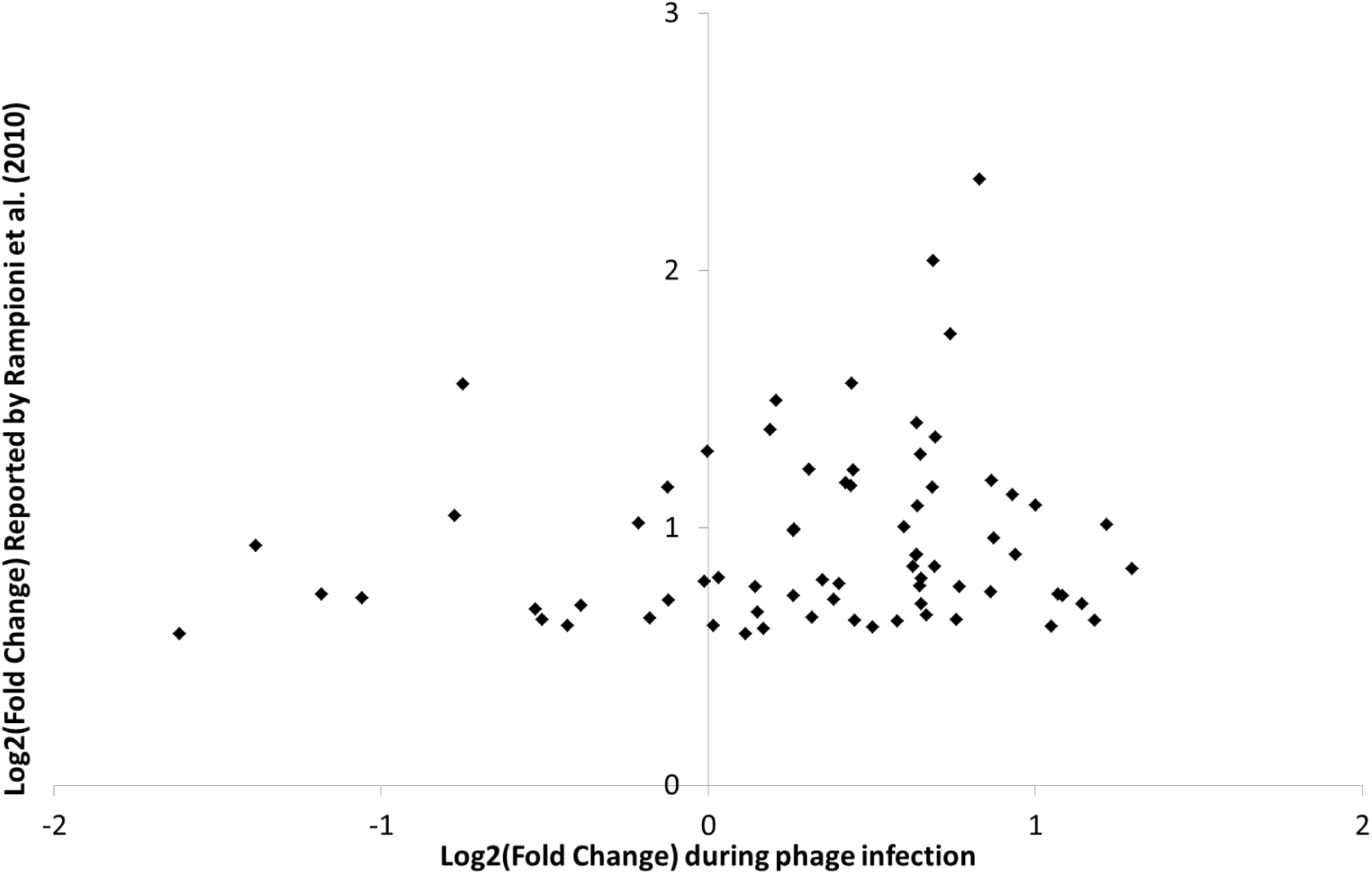
Gene features differentially expressed by PQS. The gene features found by *Rampioni et al*. (*2010*) to be upregulated in a ApqsAmutant defective in the PQS system are also inversely differentially expressed after infection by *Pseudomonas* phages, indicating that phage infection induces a PQS mediated response.

This model presents a fascinating example of the power of kin selection to drive the evolution of traits with no direct benefit to the organism that the traits are active in. Indeed, no matter how strong the production of the PQS signal is in a *Pseudomonas* cell, by functioning as a toxin the system would seem unlikely to save an infected cell and allow it to recover as a colony forming unit. Similarly, this system would not seem to provide much benefit to the cell's mostly clonal sisters in its micro-colony, which would still stand a high chance of being killed anyway in spite of the chemical warning to shut down metabolism and produce the oxidatively toxic PQS signal to harm infected cells unable to transcriptionally respond. Instead, the most significant benefit of maintaining a PQS system able to warn a micro-colony of impending phage attack would be to neighboring micro-colonies. In addition to being upregulated on a transcriptional level, we have previously found that PQS metabolites, as well as precursors, have been shown to increase in abundance as determined by whole cell mass spectrometry. This has been found within cells infected by most but not all phages (*De Smet et al., 2016*). This suggests that some of the phages with successful infections that were examined, both here and by *De Smet et al*. (*2016*), may possess phage-encoded inhibitors that prevent PQS production on a post-transcriptional level while others may have mechanisms causing the host cell to be insensitive to its effect.

Amino acid catabolism is upregulated in response to phage infection Phage infection appears to upregulate the production of transcripts for PA2441-2446, which are involved in the glycine cleavage system. This operon is under the control of GcsR (PA2449), which is not differentially expressed on a transcriptional level. However, GcsR activity does indeed appear to be induced by the host during phage infection (See Figure 5). The glycine cleavage system forms a metabolic pathway that converts glycine into pyruvate to power central metabolism, which must be tightly regulated (*Kikuchi, 1973*; *Sarwar et al., 2016*). The host may be post-transcriptionally activating GcsR so as to specifically over-regulate the transcription of the glycine cleavage system during infection, above the level that was previously expressed under phage-free conditions in the experimental flask.

**Figure 5.**
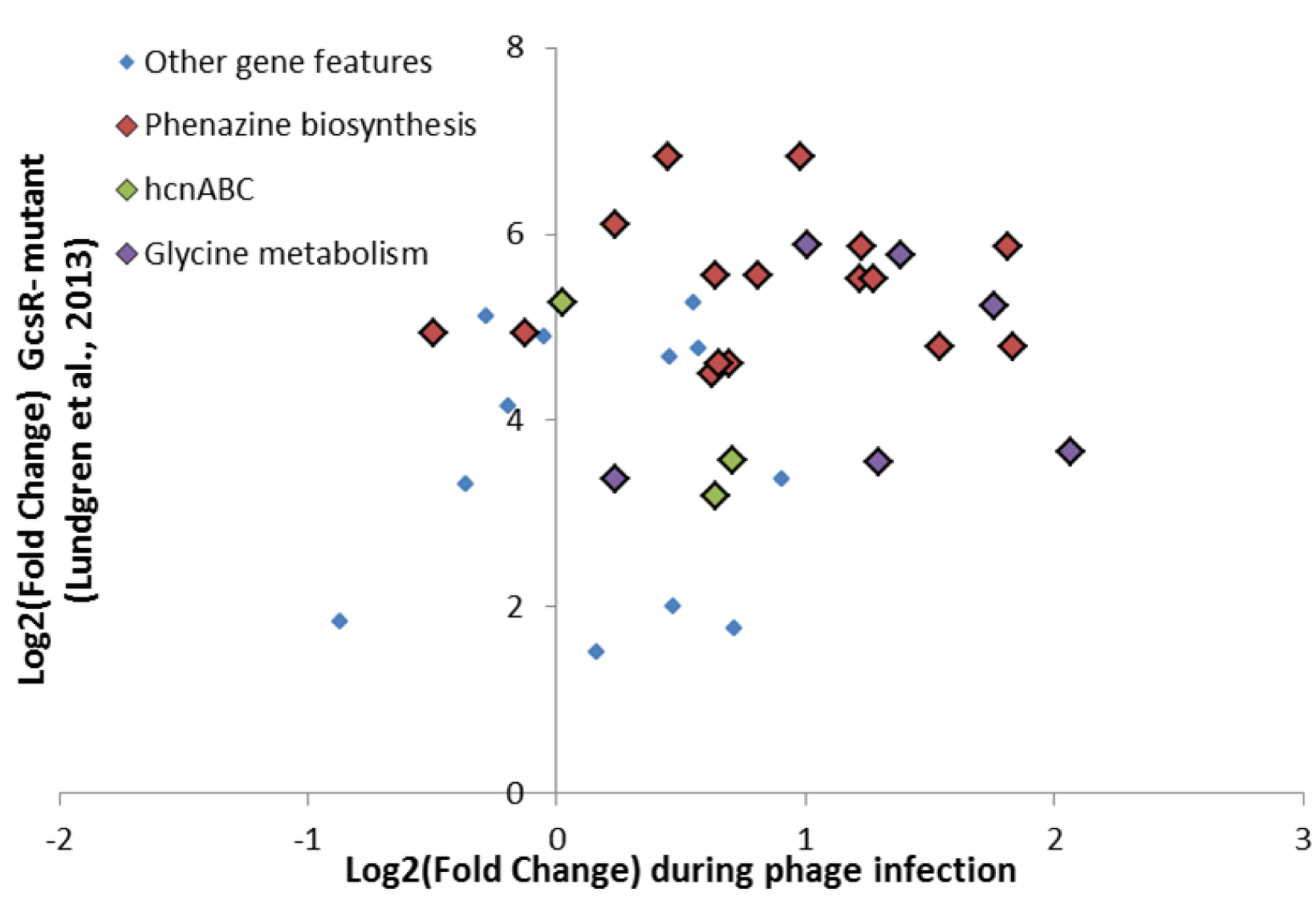
Gene features differentially expressed by GcsR. The gene features found by *Lundgren et al*. (*2013*) to be upregulated in a AGcsRmutant defective in the Glycine cleavage system are also inversely differentially expressed after infection by *Pseudomonas* phages, indicating that phage infection induces a GcsR mediated response.

Another four coding sequences, MaiA/FahA/HmgA and hpd, related to tyrosine/phenylalanine and leucine catabolism are also upregulated by the host. According to the Pseudomonas Genome Database, this pathway is predicted to encode for a 4-hydroxyphenylpyruvate dioxygenase (hpd) homogentisate-1,2-dioxygenase (HmgA), maleylacetoacetate isomerase (MaiA), and fumarylacetoacetase (FahA), which process the p-hydroxyphenylpyruvate precursor for tyrosine and phenylalanine into acetoacetate and fumarate (*Winsor et al., 2016*). However, this activation does not appear to be similar to the ordinary transcriptional response to additional phenylalanine and tyrosine (*Palmer et al., 2010*). The stress response similarly upregulates the *liuRABCDE* cluster, which converts leucine into acetoacetate for injection into central metabolism (*Aguilar et al., 2006*).

These genes involved in amino acid catabolism appear to be used as part of a ‘Scorched Earth’ strategy for reducing the vigor of infection, disrupting phage propagation without necessarily excluding it in a fashion first postulated by *Hyman and Abedon (2010)*. As *De Smet et al*. (*2016*) showed, *P. aeruginosa* phages generally appear to consistently deplete amino acid metabolites indicating that their abundance may be a major limiting factor in the success of phage infections. Indeed, this aspect of phage infection may be a particularly vulnerable target for the dying host to attack. Successful abortive infection may not require special or exotic Toxin/Anti-toxin systems (*Forde and Fitzgerald, 1999*; *Chopin et al., 2005*; *Snyder, 1995*), but could potentially simply involve the host disrupting the metabolic health of the cell in a way that the infected cell would have a hard time recovering from.

### Conservation of the PAO1 host response in PAK

Data from RNA-Seq experiments that had been previously published for infections of PAO1 by ΦKZ (*Ceyssens et al., 2014*) and infections of *P. aeruginosa* PAK by PAK_P3 (*Chevallereau et al., 2016*) and PAK_P4 (*Blasdel et al., 2017*) were also reanalyzed. With the host-mediated transcriptional stress response used by *P. aeruginosa* PAO1 during infection by various phages identified, it then became possible to assess whether this same stress response was used by *P. aeruginosa* PAK during infection by phages PAK_P3 and PAK_P4 (*Blasdel et al., 2017*). To accomplish this, coding sequences in *P. aeruginosa* PAK that were orthologous to the genes used by *P. aeruginosa* PAO1 to respond to phage infection were first identified by the Best Bi-directional Hit (BBH) method. Of the *P. aeruginosa* PAO1 coding sequences that are differentially expressed in response to phage infection, just over half could be identified in *P. aeruginosa* PAK and shown to be differentially expressed during infection by phages PAK_P3 and PAK_P4. Of the remaining half, 2/3s were not differentially expressed and 1/3 had no BBH (See Figure 6). Specifically, the PQS system was found to be unregulated in PAK while CrcZ and Hfq were found to be downregulated, suggesting that both may function in conserved ways (See Appendix Box 1). At the same time, coding sequences involved in amino acid catabolism were not found to be differentially expressed.

**Figure 6.**
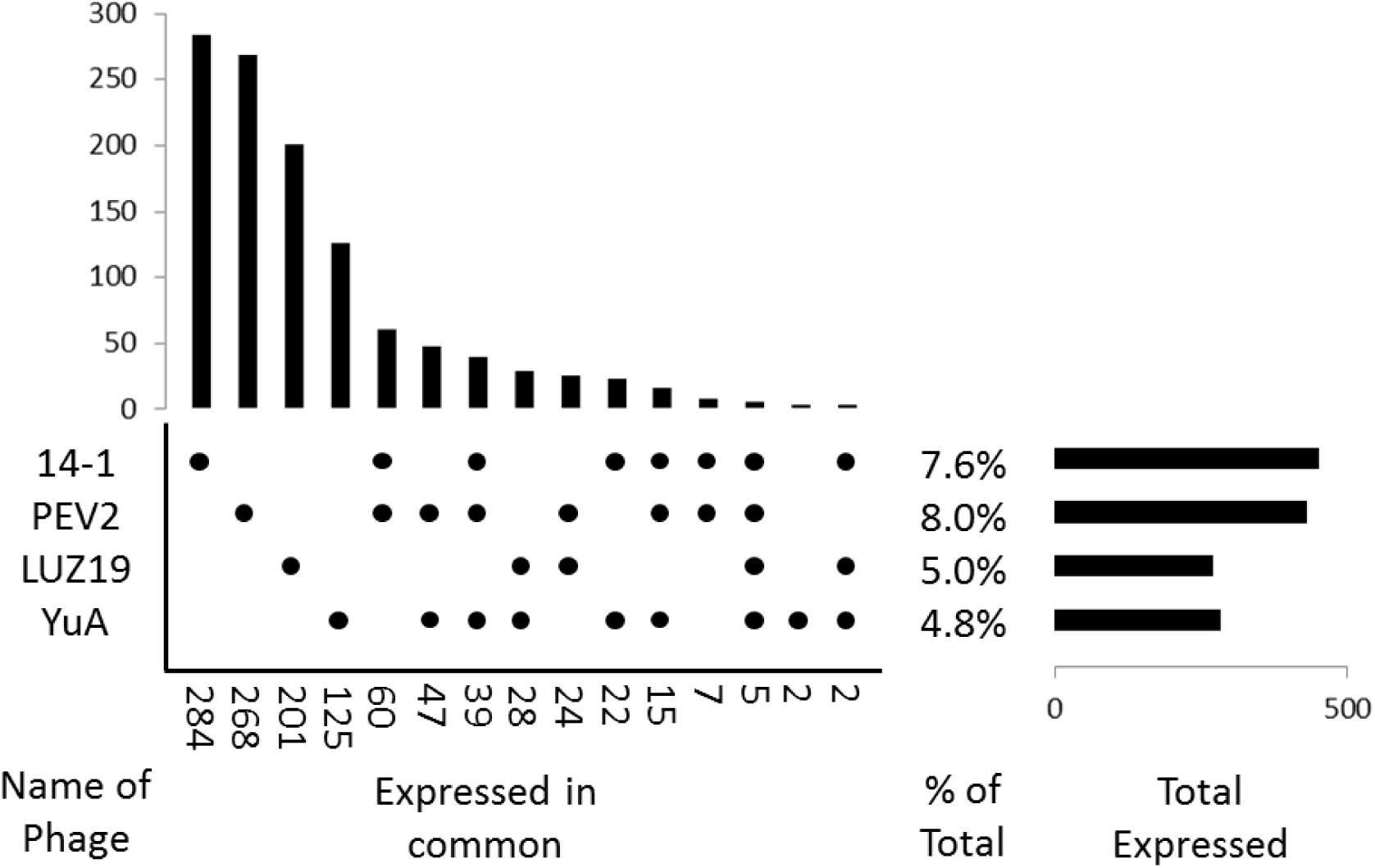
Coding sequences of the *P. aeruginosa* PAO1 stress response identified in *P. aeruginosa* PAK. Nearly half of the PAK coding sequences orthologous to genes used in the PAO1 phage stress response as identified by BBH were also differentially expressed during infection by PAK_P3 and PAK_P4(*Blasdel et al., 2017*).

**Figure 7.**
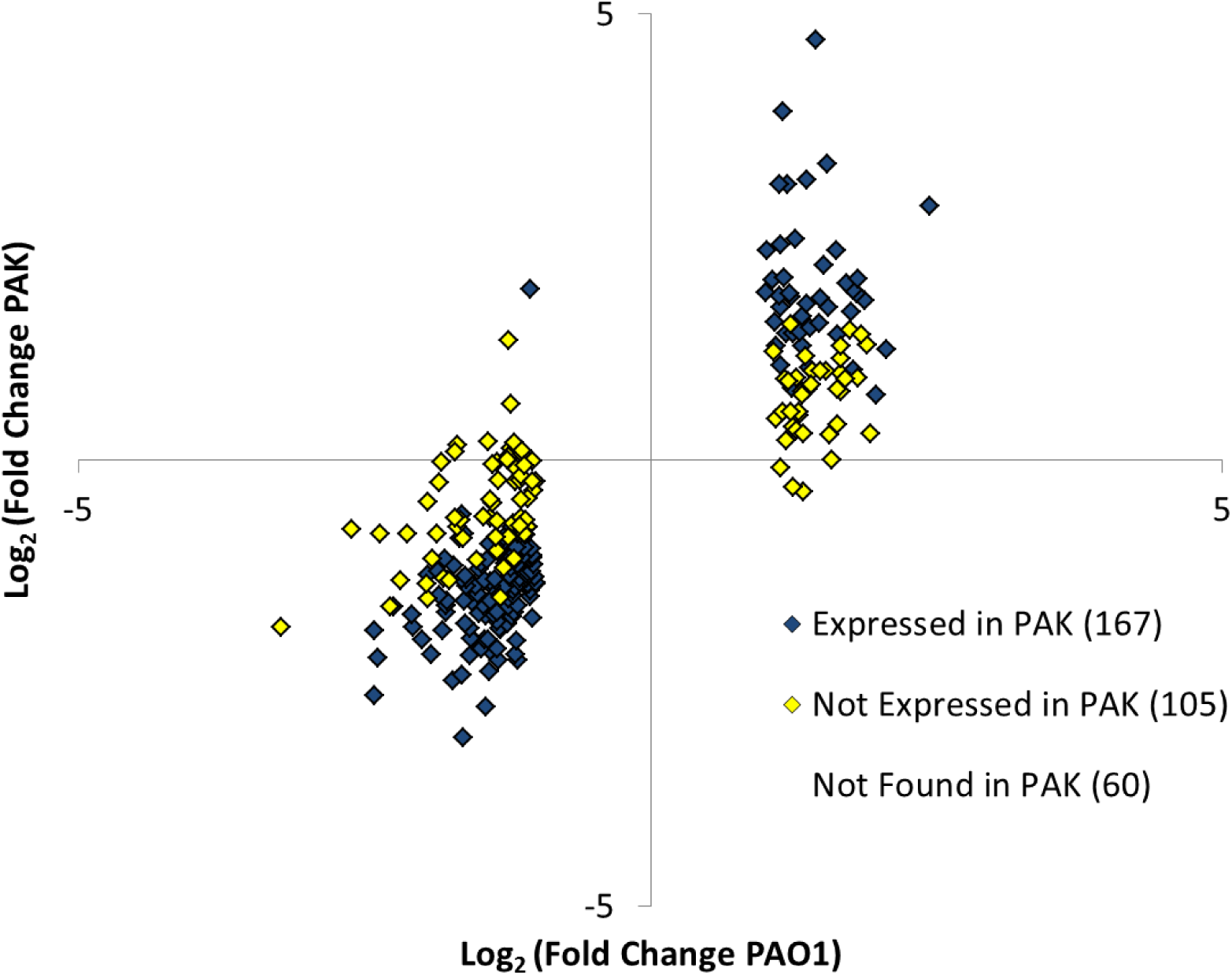
Number of host gene features that are differentially expressed during infections by each combination of phage(s) excluding the host response. We find that each phage has a mostly unique impact on host transcription.

**Figure 8.**
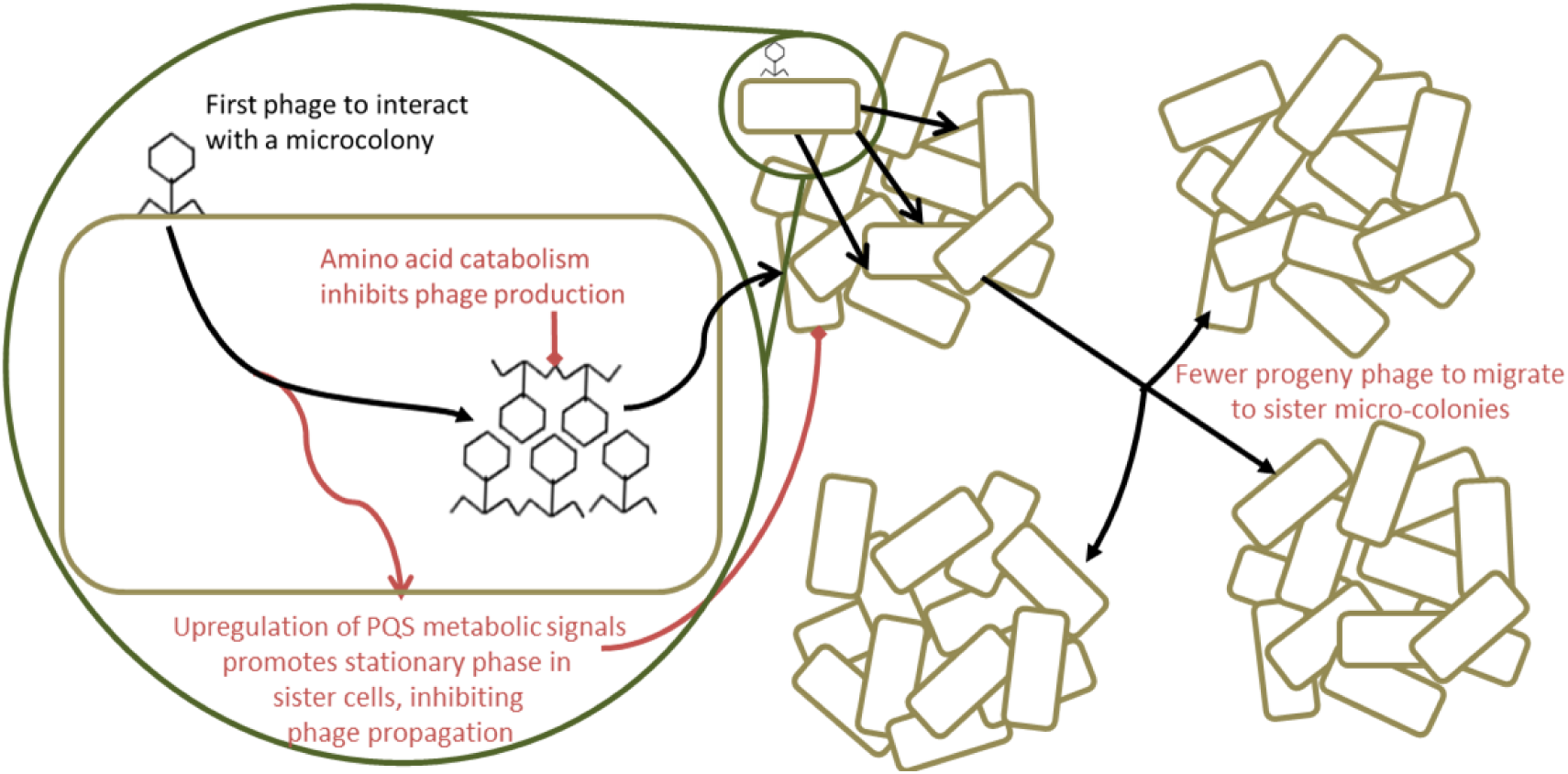
Proposed models for adaptive phage resistance. We propose that the PQS quorum sensing system promotes bacterial fitness in the presence of phage, as was found by *Moreau et al*. (*2017*) by warning sister cells to enter stationary phase, making them less susceptible to phage infection and suppressing phage propagation. Similarly, we propose that the non-adaptive upregulation of amino acid catabolism that we observe could function to inhibit phage propagation.

### Phage-mediated differential expression of the host genome

Having defined the host-mediated stress response to phage infection, we can set it aside to look specifically at each of the other gene features differentially expressed during each of the individual phage infections. Remarkably, while each phage manipulates between 4.8% and 8.0% of host gene features (Log2 fold change >1.3, adj pvalue <0.05), 20% are manipulated by at least one of the four phages, indicating a wide diversity of strategies for co-opting the functions of host gene features. Indeed, within this 20%, 78% are manipulated by just one phage. Additionally, out of the total number of gene features that are significantly expressed during more than one phage infection, 56% are significantly differentially expressed in the analysis designed to highlight the host-mediated response.

We have found that all but one of the *Pseudomonas aeruginosa* phages studied here manipulate expression of the host genome during infection, while simultaneously allowing the host to transcriptionally respond to the phage. Therefore we observe a mixture of two kinds of differential expression of host genes that could each be described as phage-mediated: (i) activation of various kinds of stress responses to metabolic and environmental stresses that the phage imposes (indirect phage manipulation) and (ii) induction of differential expression of targeted parts of the host genome through phage-borne molecular mechanisms (direct phage manipulation). At the same time, the implications of this finding are more concretely interpretable with an understanding of phage biology that centers on the phage-infected cell rather than on the inert viral particle, as in the virocell model (*Forterre, 2011*, *2013*). Indeed, this shift in perspective, which would see the phage-infected cell as a living entity that is propagated by its particles, would allow us to posit that the genome of host origin within the infected *P. aeruginosa* cell can function as a second, auxiliary, phage genome for at least many phages.

### Phage PhiKZ inhibits all differential expression of host transcripts

Phage ΦKZ is an unusually large (280kb) myovirus infecting *P. aeruginosa* that was previously understood to encode 306 coding sequences (*Mesyanzhinov et al., 2002*), later updated to 369 based on RNA-Seq data (*Ceyssens et al., 2014*). Recently, *Ceyssens et al. (2014*) demonstrated that ΦKZ can infect its host, expressing three separate phases of transcription, completely independently of the action of the host RNA polymerase. Indeed, they found that ΦKZ proceeds in the presence of high concentrations of rifampin, which acts as a powerful inhibitor of all host transcription by inactivating the host RNA polymerase. This independence was then concretely demonstrated by *Yakunina et al*. (*2015*) to result from the action of two enzymes with RNA polymerase activity, one which is packaged into the virion and co-injected with ΦKZ DNA to power early transcription (vRNAP), while the other is encoded by middle transcripts and powers late transcription (nvRNAP).

Unlike each of the other phages examined here (See Supplementary Table 1), ΦKZ infection does not appear to either prompt the host-mediated response to phage infection or mediate phage-specific differential expression of the host genome. Indeed, the genes identified as being upregulated as part of the host-mediated response to other phages are not significantly differentially expressed and there do not appear to be any host genes that are upregulated uniquely by ΦKZ. Similarly, the differential degradation of host transcripts by either entity within the cell appears to be completely disrupted by 10 minutes into infection. The only exception to this is the transcriptional activation of a Pf1-like prophage, which appears to recruit the ΦKZ early transcriptional apparatus. This suggests that ΦKZ encodes for an immediate-early inhibitor that prevents host transcription in addition to gp37, which is known to inhibit the host RNA degradosome (*Van den Bossche et al., 2016*). Thus ΦKZ, like the *Tevenvirinae* are already known to do (*Uzan, 2009*; *Doron et al., 2016*), may have made an evolutionary tradeoff whereupon they make themselves insensitive to any adaptive phage defenses that the host may use, but in exchange also render all of the other stress responses and genetic information encoded by the host useless to them. ΦKZ is notably one of the two largest phages assessed here, suggesting that larger phages like it and the *Tevenvirinae*, that encode a large number of genes could potentially adapt more easily to not requiring the information encoded by the host.

### Unsuccessful induction of prophage transcription

Of the seven phages examined here, five induce the transcription of a prophage element within the *P. aeruginosa* PAO1 or *P. aeruginosa* PAK hosts. Two of these five prophage inductions appear to conspicuously fail as only a third of each prophage element's coding sequences is differentially expressed before the end of infection (i.e. inLUZ19and PEV2-infected cells). Of the remaining three, ΦKZ as been previously demonstrated to fail to fully mobilize the P1-like prophage element encoded by PAO1 as *Ceyssens et al*. (*2014*) showed that the element does not replicate its own genome. Similarly, as was reported by *Chevallereau et al*. (*2016*) and *Blasdel et al*. (*2017*) respectively, PAK_P3 and PAK_P4 both appear to fail to fully mobilize the P2-like prophage element encoded by PAK. Indeed, they have shown that in both cases the upregulation of the prophage element relative to other host transcripts does not outpace the more global replacement of host transcripts with phage transcripts leading them to be ultimately downregulated relative to the total transcript population in the infected cell (See Table 2).

**Table 2.**
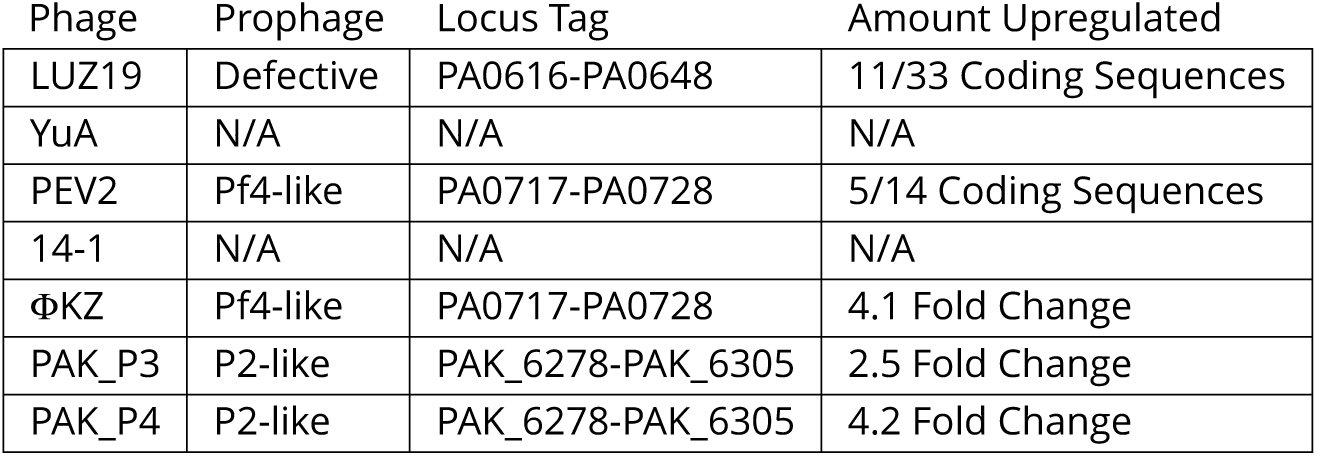
Prophage elements induced during infections by various phages. Of the seven phage infections examined in this manuscript, three promote the transcription of a prophage within *P. aeruginosa* PAO1 or PAK, two promote the transcription of part of one prophage, and two do not promote prophage transcription.

| Phage | Prophage | Locus Tag | Amount Upregulated |
| --- | --- | --- | --- |
| LUZ19 | Defective | PA0616-PA0648 | 11/33 Coding Sequences |
| YuA | N/A | N/A | N/A |
| PEV2 | Pf4-like | PA0717-PA0728 | 5/14 Coding Sequences |
| 14-1 | N/A | N/A | N/A |
| ΦKZ | Pf4-like | PA0717-PA0728 | 4.1 Fold Change |
| PAK_P3 | P2-like | PAK_6278-PAK_6305 | 2.5 Fold Change |
| PAK_P4 | P2-like | PAK_6278-PAK_6305 | 4.2 Fold Change |

## Conclusions

While it has been long established that T4 both shuts off the transcription and translation of all host gene features (*Koerner and Snustad, 1979*; *Black et al., 1983*; *Kutter et al., 1994*), and that T5 so thoroughly degrades host DNA that it is implausible that any is transcribed during infection, it has previously been unclear if this exclusion of host transcription is universal or even common. Indeed, *Schachtele et al. (1972)* demonstrated through radioactive experiments that most transcripts produced during Φ29 infection of *Bacillus subtilis* originate from the host, suggesting that they may play a role in infection without being able to indicate what that role might be. There are clear benefits to the phage in shutting off the host's ability to mount a stress response on a transcriptional level. By leaving host mechanisms able to detect and respond to infection through adaptive processes, a phage would leave itself vulnerable to unpredictable host-mediated interference. However, not only would this inactivation inhibit host defense responses that rely on gene expression, but it would also serve to exclude the expression of other competing selfish genetic elements within the cell that might interfere with productive infection such as prophages, competing phage particles, and plasmids. At the same time, the potential benefits to allowing the host to transcriptionally respond to stress are also significant. Phages that allow the host transcriptional system to remain intact at least on some level could manipulate host stress responses directly with phage machinery, as well as indirectly through phage-induced stress, and thus adapt the cell to phage specific conditions. We have observed that four out of five examined phages that infect *P. aeruginosa* PAO1 prompt the activation of our proposed Bacterial Adaptive Phage Response (BAPR) systems while specifically mediating differential expression of the host genome.

### PAO1 shows a BAPR response which potentially limits phage infection

The two community-focused systems for defense against phage predation that we present, PQS-mediated quorum sensing and amino-acid catabolism, appear to be examples of a new class of phage defenses. While both the PAO1 and PAK strains examined in this report lack many of the intrinsic and acquired defenses against phage infection known to be present in *Pseudomonas* spp., such as CRISPR-Cas, the potential for a communal response signaling phage predation could go a long way towards explaining the success of these systems in strains that have them. Constitutive expression of the CRISPR-Cas system in PA14 has been previously shown to carry significant fitness costs (*Westra et al., 2015*), possibly stemming from autoimmunity (*Vercoe et al., 2013*), and a molecular forewarning of impending phage infection could activate CRISPR-Cas and other known systems for promoting phage resistance. Indeed *Høyland-Kroghsbo et al. (2016)* have previously shown expression of CRISPR-Cas transcripts to be induced by the LasR quorum sensing system in *P. seudomonas* PA14, and the ability of dying cells to send a final warning could help explain the function of that system.

Notably, overproduction of the PQS signal has also long been observed to promote autolysis, likely through the induction of prophages, in *P. aeruginosa* (*D’Argenio et al., 2002*) as well as P putida (*Fernández-Piñar et al., 2011*). Indeed, the connection between lytic phage infection, intercellular signaling, and prophage induction could reflect *Pseudomonas* prophages sensing the host's call to arms and using it as a signal to escape. We have observed three prophage elements beginning, and failing, to autoinduce in response to phage infection which may be in part due to PQS warning. This connection could also help explain the notorious difficulty of isolating and maintaining pure stocks of phages when clinical or environmental strains, which often carry active prophages and have not been adapted to laboratory conditions, are used as hosts.

We have also found what may be considered as a ‘Scorched Earth’ strategy for reducing the vigor of phage infection, whereby an infected cell adaptively disrupts its own metabolism in such a way as to impair the success of phage infection. This would serve as a practical example for systems of ‘Reduced Infection Vigor’ that have been previously postulated by *Hyman and Abedon (2010)*.

## Methods and Materials

### Strains and growth conditions

RNA sequencing of phage-infected cells was performed as described by *Chevallereau et al*. (*2016*) and *Blasdel et al*. (*2017*). The *Pseudomonas aeruginosa* strain PAO1 was used as a host for phages LUZ19, YuA, PEV2,14-1, and ΦKZ. Cultures were grown to an OD_600_0.3 (1.2 × 10^8^ / mL) in Lysogeny broth (10 g/L Tryptone, 5 g/L Yeast extract, 5 g/L NaCl) and infected at a multiplicity of infection of 50 to ensure synchronous infection. *P. aeruginosa* strain PAK was used as a host for phages PAK_P3 and PAK_P4.

### Whole transcriptome sequencing

RNA-Sequencing was performed on four phage-negative controls taken immediately before infection. Samples were also taken from synchronized infections by phages LUZ19, YuA, PEV2, and 14-1 to represent early, middle and late transcription as described by *Blasdel et al. (2018*, *2017)*. Briefly, synchronicity was controlled for by ensuring that fewer than 5% of host cells still survived before the early time point was taken. At time points after the addition of phage chosen to reflect each phase of infection, cells were immediately halted with stop solution (1%, 9% ethanol, 90% sample (*Blasdel et al., 2018*)), pelleted, and re-suspended in TRIzol to lyse cells as well as inactivate RNAses and start an organic extraction. The resulting nucleic acid was digested with TURBO DNAse to remove DNA, rRNA was removed with the Ribo-Zero kit, and stranded cDNA was prepared with the TruSeq Stranded mRNA Library Prep Kit ahead of sequencing.

### Differential expression analysis

After trimming, sequencing reads were aligned separately to both the phage and host genomes with the CLC Genomics workbench v7.5.1 (QIAGEN Bioinformatics, Aarhus, Denmark). Sequencing statistics and results are summarized in Supplementary Table 2. Alignments were then summarized into count tables of Total Gene Reads that map to phage or host gene features respectively. Each statistical comparison presented was performed using the DESeq2 (*Love et al., 2014*) R/Bioconductor package to normalize host transcript populations to host transcript populations, before testing for differential expression as described by *Blasdel et al*. (*2017*).

The core analysis compares directional host total gene read counts for four phage negative controls sampled at a point immediately before infection to samples representing the synchronized late infections of LUZ19, YuA, PEV2 and 14-1. The DESeq2 test for the statistical significance of differential expression between the negative controls and phage-infected samples here is used to specifically highlight gene features that are differentially expressed in common between the different phage infections (See Figure 2). As the four phages used have radically different metabolisms, genome contents, and methods for coopting host mechanisms, they are unlikely to all mediate differential expression of any given host gene in the same way. Thus, any differential expression of host genes that is common to all of the independent infections is likely to be mediated by the one biological entity all of the infections have in common, the host.

#### Normalization of reads between conditions

Given that there are two biological entities within the infected cell whose transcripts are sequenced together, and which change in total abundance relative to each other, meaningfully assessing differential expression for host genes becomes non-intuitively complex. Indeed, we observe a global depletion of all host transcripts relative to the total (phage and host) transcript population present in the cell due to the influx of phage transcripts (See Figure 1). Therefore, two approaches could theoretically be used to measure the change in the abundance of a host transcript during a phage infection. We could measure changes (1) relative to either only other host transcripts or (2) relative to the total transcript population in the cell, including new phage transcripts. We have here adopted the first, asking which host transcripts are up or down regulated relative to other host transcripts by normalizing host transcripts before infection to host transcripts after infection. This has the effect of artificially enriching host reads in the phage infected condition by normalizing away the influx of phage transcripts, so as to compare it to the uninfected condition as if those phage transcripts were not present. The answer to this question focuses on how the host transcripts are being being targeted for either increased transcription or decreased degradation by at least one of the two biological entities in the cell, by ignoring the replacement of host transcripts with phage transcripts.

Analyzing the change in a transcript's abundance relative to the total transcript population could not be addressed analysis of multiple phage infections, since depletion of host transcripts relative to the total happens at different rates during different phage infections. This effectively prevents us from elucidating a transcript's differential ability to compete for ribosome activity through changes relative abundance. For example, as was reported for prophage transcripts by *Blasdel et al*. (*2017*) during phage infection, a transcript could be targeted for increased transcription even while the stronger degradation of all non-phage transcripts reduces its abundance relative to the total transcript population in the cell.

Indeed, the differential expression results that we report will not reflect a transcript's differential ability to compete for ribosome activity through changes relative abundance. To answer this second question, we would need to ask which host transcripts are up or down regulated relative to the total transcript population in the cell, including both phage and host transcripts. F

It is important, however, that the changes in the relative abundance of host transcripts relative to other host transcripts that we are measuring in this manuscript happen independently of the global depletion of host transcripts relative to the total transcript population. Thus, even though the depletion of host transcripts relative to the total happens at different rates during different phage infections, we are still able to meaningfully test host genes for differential expression in the comparison between our phage negative controls and the combined phage-infected condition that forms our core analysis.

#### Comparisons to other data sets

The gene features identified by this analysis as being differentially expressed in common between each infection (See Figure 2) were then highlighted in data published for ΦKZ (*Ceyssens et al., 2014*) as well as each phage examined (See Figure 3-Figure supplement 1). Orthologues to gene features in the PAO1 host response to phage infection were identified in the PAK strain by the Bi-directional Best Hit method. The differential expression identified in the core analysis was also compared to published data representing the differential expression between wild-type PAO1 and a AHfqmutant (*Sonnleitner et al., 2006*), a ΔGcsR-mutant (*Lundgren et al., 2013*), as well as a APqsA-mutant (*Rampioni et al., 2010*). Neither the three wild-type negative control conditions nor the three mutant conditions used by each report can be directly compared to any of the conditions used here due to the differences in expression that can be expected in bacteria growing in different labs. However, this analysis compares only the change in expression due to phage infection to the change in expression due to the absence of each gene.

## Acknowledgments

We would like to gratefully acknowledge the thoughtful critique and thoughts of Dr. Harold Brussow. We would also like to thank Dr. Kasia Dani-Wtodarczyk and Maarten Boon for technical assistance in collecting RNA samples from synchronous infections and Dr. Jeroen De Smet for his assistance in interpreting his metabolomics data. B.G.B acknowledges the Onderzoeksfonds K.U. Leuven for granting a postdoctoral fellowship (3E160483). This research was also supported by grant G.0323.09 from the FWO and the KULeuven project GOA ‘Phage Biosystems’.

## Appendix 1 Hfq transcripts are depleted in response to phage infection

The *Pseudomonas* Hfq peptide has been demonstrated to act as an RNA chaperone for mRNAs and sRNAs, positively and negatively modulating both abundance and function of each RNA species (*Vogel and Luisi, 2011*; *Geissmann and Touati, 2004; Valenntini et al., 2011*).

We have found that the host response to phage infection appears to differentially express the same gene features that *Sonnleitner et al*. (*2006*) found to be differentially expressed in an ΔHfq-mutant when compared to a wild type control (See Figure 1), suggesting an absense of Hfq activity. Indeed, 75% of genes that are upregulated in the ΔHfq-mutant are also upregulated during phage infection, while both conditions only significantly modulate 14% and 5.5% of the host gene features respectively. While the transcript encoding for Hfq is downregulated during infection by each Pseudomonas phage examined, including PAK_P3 and PAK_P4 infecting the PAK strain, the CrcZ sRNA is also just as universally rapidly depleted. CrcZ was recently demonstrated by *Sonnleitner and Bläsi (2014)* to titrate Hfq *in vitro* abrogating Hfq-mediated translational repression. Indeed, CrcZ appears to decoy for Hfq, competitively inhibiting each of its mRNA dependent roles (*Sonnleitner and Bläsi, 2014*).

**Appendix 1 Figure 1.**
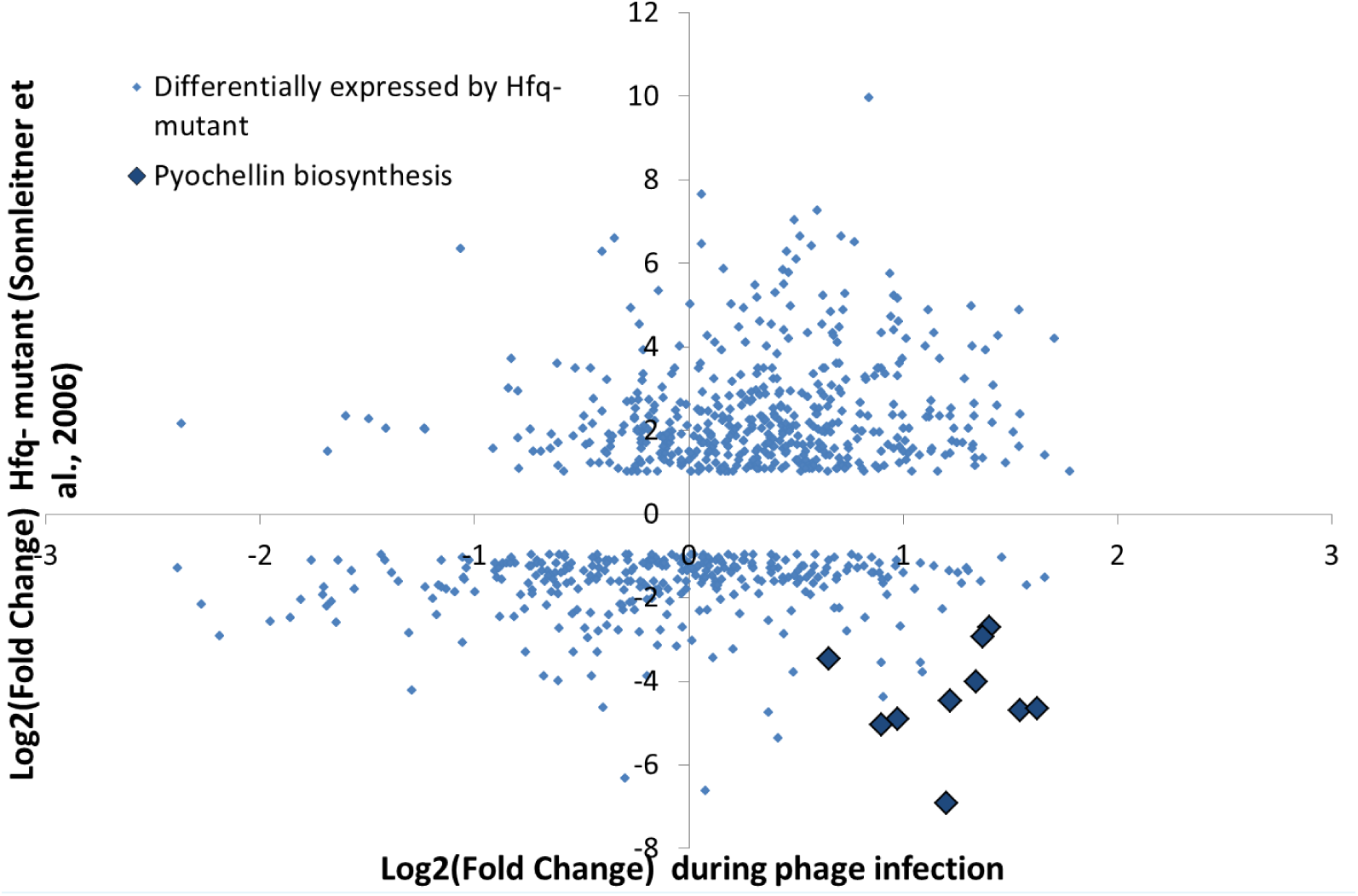
Gene features differentially expressed by Hfq The gene features found by *Sonnleitner et al*. (*2006*) to be differentially expressed in a ΔHfq-mutant are also inversely differentially expressed after infection by *Pseudomonas* phages, indicating that phage stress infection induces Hfq activity. CrcZ is not reported as differentially expressed by *Sonnleitner et al*. (*2006*).

Within the stress response we observe the downregulation of CrcZ, which would presumably liberate Hfq activity titration mediated by this sRNA. However, we also observe the downregulation of Hfq transcripts, and the total impacts of these two effects on Hfq-regulated transcripts is similar to a ΔHfq-mutation. At the same time, microbiological evidence suggests that one of the many functions of Hfq may be useful as a phage defense, indeed phage 14-1 was notably observed to produce three fold more plaques on a ΔHfq-mutant as it does on its WT primary host. Remarkably, Hfq was first described and named as the essential component of the Host Factor fraction required for plus strand synthesis from RNA phage Q*β* (*Franze de Fernandez et al., 1968*), suggesting that Hfq activity may have a complex and multifactorial relationship to phage infection.

### Host stress response upregulates pyochelin biosynthesis

One puzzling feature of the host response is its upregulation of the pyochelin iron uptake pathway (pchABCDEFGR), contrary to what could be expected given the inhibition of Hfq activity that is observed. Pyochelin is one of two primary siderophores secreted by *P aeruginosa*, the other being pyoverdine, to chelate and adsorb iron (*Braud et al., 2009*). Siderophores are postulated to be involved in phage specific manipulations of the host during PAK_P4 infection of *P. aeruginosa* PAK (*Blasdel et al., 2017*). However, we are at a loss to explain how or why pyoverdine is more weakly upregulated by the host.

### Host stress response upregulates an efflux pump system

One of the major PAO1 efflux pump systems (MexCD/OprJ) appears to be transcriptionally upregulated in response to phage infection. The MexCD/OprJ system has been demonstrated to effect the extrusion of a large number of critical antibiotics (*Gotoh et al., 1998*; *Masuda et al., 2000*). While it is unclear why the host is responding to phage infection with this efflux pump system, it may reflect an attempt to produce abortive infection by extruding an unknown metabolite that could be vital to some phage infections.

**Figure 3-Figure supplement 1.**
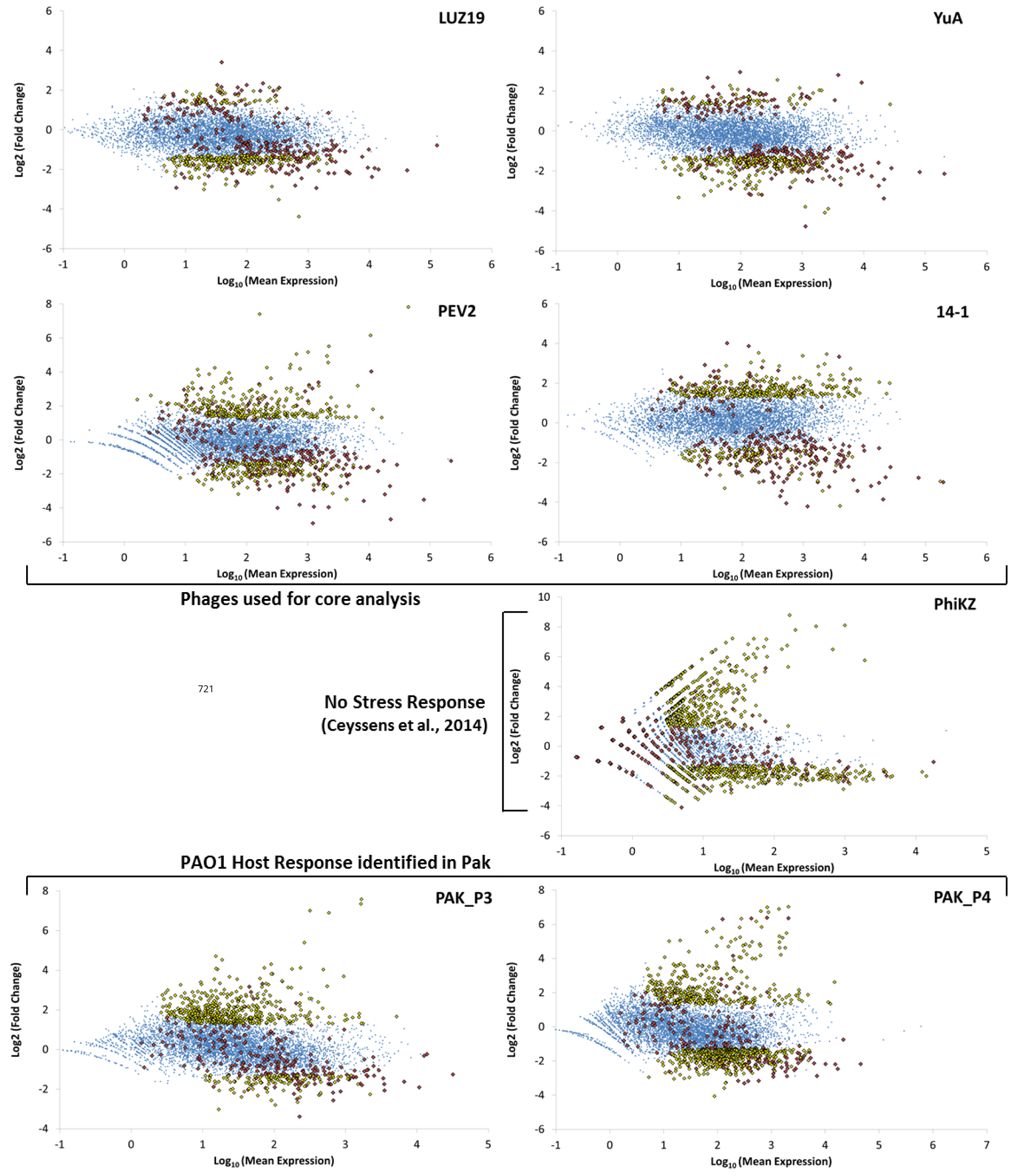
The differential expression of host genes during infection by each phage are examined here individually. Gene features identified as differentially expressed by the putative host-mediated stress response are highlighted in each individual phage infection as red, while gene features that are differentially expressed by each phage but not the common host response are highlighted as yellow. The differential expression of the putative host-mediated stress response appears to be weaker in LUZ19 than in other phage, though this could be explained by the activity of LUZ19 gp25.1, a T7 gp2-like transcriptional inhibitor (*Klimuk et al., 2013)* in middle infection.

